# Multi-modal single-cell and whole-genome sequencing of minute, frozen specimens to propel clinical applications

**DOI:** 10.1101/2022.02.13.480272

**Authors:** Yiping Wang, Joy Linyue Fan, Johannes C. Melms, Amit Dipak Amin, Yohanna Georgis, Patricia Ho, Somnath Tagore, Gabriel Abril-Rodríguez, Jana Biermann, Matan Hofree, Lindsay Caprio, Simon Berhe, Shaheer A. Khan, Brian S. Henick, Antoni Ribas, Alison M. Taylor, Gary K. Schwartz, Richard D. Carvajal, Elham Azizi, Benjamin Izar

**Author notes:** equal contribution (Yiping Wang, Joy Linyue Fan, Johannes C. Melms).

## Abstract

Single-cell genomics are enabling technologies, but their broad clinical application remains challenging. We report an easily adaptable approach for single-cell transcriptome and T cell receptor (TCR)-sequencing, and matched whole-genome sequencing from tiny, frozen clinical specimens. We achieve similar quality and biological outputs while reducing artifactual signals compared to data from matched fresh tissue samples. Profiling sequentially collected melanoma samples from the KEYNOTE-001 trial, we resolve cellular, genomic, and clonotype dynamics that encapsulate molecular patterns of tumor evolution during anti-PD-1 therapy. To demonstrate applicability to banked biospecimens of rare diseases, we generate a large uveal melanoma liver metastasis single-cell and matched WGS atlas, which revealed niche-specific impairment of clonal T cell expansion. This study provides a foundational framework for propelling single-cell genomics to the clinical arena.

## MAIN

Single-cell genomics, namely single-cell RNA-sequencing (scRNA-seq), has enabled significant discoveries in virtually all fields of biomedical research, including in several solid tumors, providing unique insights into tumor ecosystems^1–4^, metastasis^5,6^, and drug resistance^2,7^. Matched scRNA/TCR-seq is particularly informative for studies investigating the effect of transformative immunotherapies, such as PD-1-targeting monoclonal antibodies^8^. While these methods have the potential to inform clinical studies, their broad application has been hampered by several challenges. In particular, the need for relatively large (milligrams to grams), fresh tissue specimens and their immediate processing is incompatible with clinical workflows and prohibits streamlined multi-institutional analysis efforts. Consequently, single-cell studies to date have been conducted in relatively small, heterogeneous patient populations (e.g. variable treatment exposures), which introduce challenging biases in their interpretation. Building on recent developments that enable single-nucleus transcriptome sequencing (snRNA-seq) from frozen tissues^9,10^, we have evolved an approach to perform rapid, scalable, and high-quality single-cell transcriptome and matched TCR-seq of very small (nanograms to micrograms) clinical biopsy specimens, as well as population-matched ultra low pass whole-genome sequencing (ulp-WGS) (**Methods**).

We first performed head-to-head comparisons of scRNA/snRNA-seq of matched fresh and frozen tissues from patients with non-small cell lung cancer (NSCLC), metastatic cutaneous melanoma, and uveal melanoma using different 10X Genomics chemistries (3’, 5’v1, or 5’v2), cell sorting and RNase inhibitor protocols (**Methods**; **Extended Data Table 1**).

Overall, in snRNA-seq using 5’ chemistries, the data quality (median number of genes detected per cell) was comparable to that of tissue-matched scRNA-seq and was consistently superior compared to 3’ chemistries (**Fig. 1a-c**), while snRNA-seq protocols had a lower rate of mitochondrial reads (which increases in cells with impending death) (**Extended Data Fig. 1**). In the NSCLC comparison, for example, we recovered a median of 3,117 genes/cell using 5’v1 (with addition of RNase inhibitor) compared to 2,521 in fresh scRNA-seq (Wilcoxon rank-sum p-value 1.67e-29) and 1,392 in the best 3’ condition (Wilcoxon rank-sum p-value 0), respectively. The median fraction of mitochondrial reads accounted for 0.97% of reads in the 5’v1 snRNA-seq NSCLC specimen, compared to the matched fresh specimen with 3.94% (Wilcoxon rank-sum p-value 4.91e-280). Similar results were observed in cutaneous and uveal melanoma tissue comparisons (**Extended Data Table 2**).

**Figure 1.**
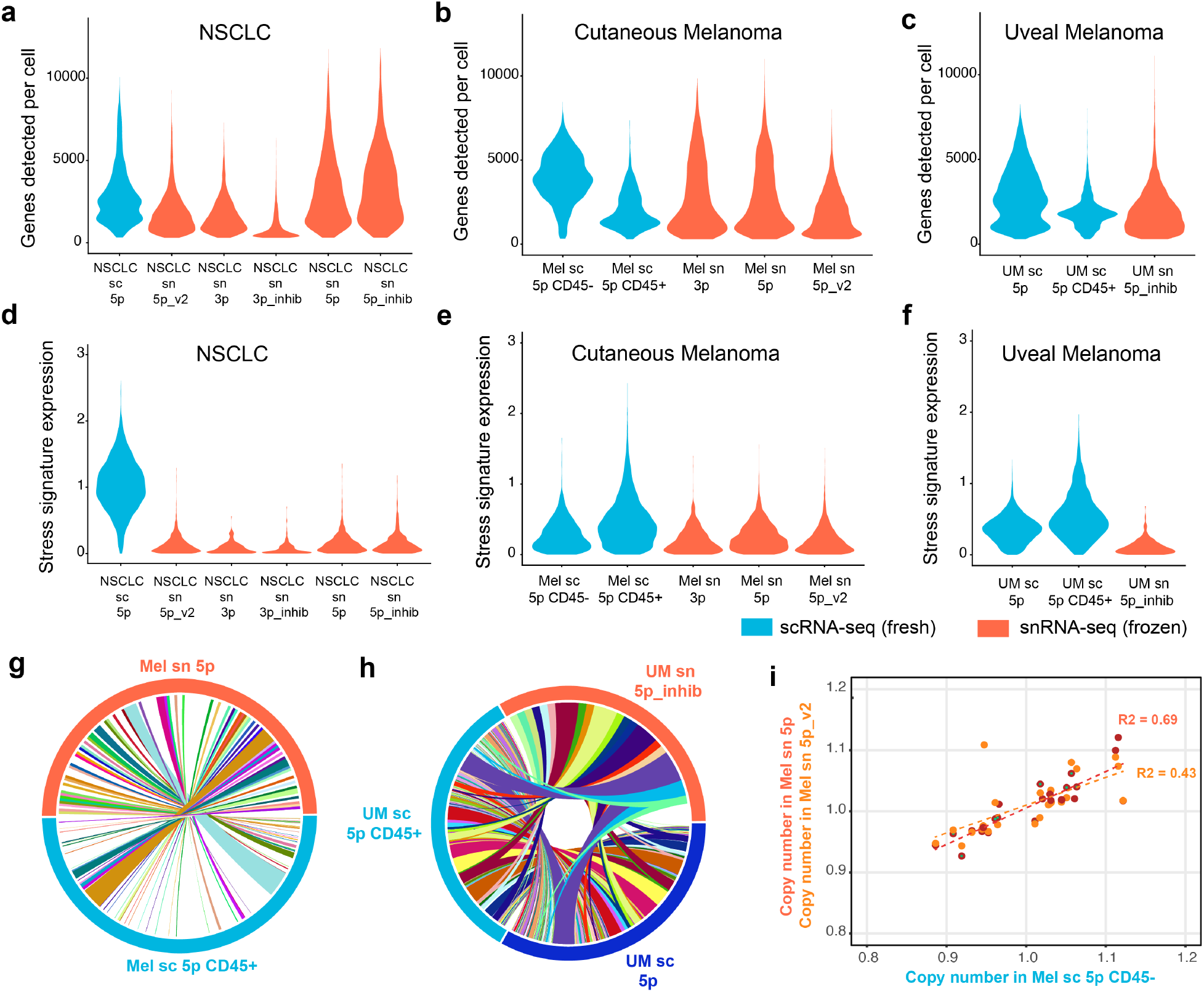
**a-c**, Violin plots of genes detected per cell in (a) non-small cell lung cancer, (b) cutaneous melanoma and (c) uveal melanoma samples. Blue plots indicate scRNA-seq from fresh tissue, and red indicates snRNA-seq from frozen tissue. 10X chemistry type and presence/absence of RNAse inhibitor is indicated on labels beneath each violin. **d-f**, Violin plots of relative expression of an artifactual stress-associated gene expression signature. Samples and experimental settings corresponding to panels (a-c) above. **g**,**h**, Circos plots of T cell receptor clonotypes in (g) cutaneous and (h) uveal melanoma, respectively. Connections indicate overlap of identical TCRs found in both fresh scRNA-seq and frozen snRNA-seq samples. **i**, Correlation of average cutaneous melanoma cell arm-level CNAs predicted by inferCNV, in a fresh CD45-scRNA-seq sample (x-axis) vs. two frozen 5p snRNA-seq protocols (y-axis).

Importantly, we noted substantial expression of an artifactual program associated with tissue processing in scRNA-seq^11^ in all three tissue comparisons, while this artifact was more lowly expressed or absent in matched snRNA-seq data (**Fig. 1d-f**). This artifactual expression manifested particularly strongly in the NSCLC comparison, was increased by fluorescence-activated cell sorting (FACS), and was more prominent in immune cells. Notably, this stress signature captures inflammatory pathways^11^, among others, and may bias the interpretation of these pathways in scRNA-seq.

In the cutaneous melanoma sample, which is rich in large chromosomal aberrations, we next inferred copy-number alterations (CNAs) and found strong agreement in average arm-level CNA (Spearman R2=0.69) between scRNA-seq and snRNA-seq protocols (**Fig. 1g**). Lastly, in 5’ chemistry sc/snRNA-seq, we were able to robustly recover TCRs matched to single-cell transcriptomes. We assessed the degree of overlap of TCR’s from frozen tissues with those recovered from the matching fresh tissue using a hypergeometric test (**Methods**). This overlap was statistically significant in all cases (cutaneous melanoma CD45+ fresh vs. 5’v1 frozen: p=1.55e-62, uveal melanoma fresh unsorted vs frozen with inhibitor: p=0.00052, uveal melanoma fresh CD45+ vs 5’v1 frozen: p=0.0018). Fractional clonotype sizes (**Fig. 1h,i**) and TCR diversity, as assessed by the Gini coefficient (**Extended Data Table 3)**, were also comparable between fresh and frozen samples, as well as in the sequential samples from an anti-PD1 therapy patient (**Extended Data Fig. 2**).

Despite procedural and technical differences, matched sc/snRNA-seq showed good mixing of cell types (determined using the LISI score^12^, **Methods**) following batch correction (**Methods**) indicating preservation of global transcriptional outputs (**Extended Data Fig. 3,4**). Accordingly, we identify comparable cell type diversity (estimated using the Shannon index, **Extended Data Fig. 5a,b**) across comparisons. Notably, cell type composition was highly consistent among different snRNA-seq runs within the same specimen, indicating that these methods are highly robust (**Extended Data Fig. 5a**). Comparing sc/snRNA-seq, we noted one important outlier in cellular composition: there was low recovery of cancer cells in the NSCLC sample profiled using scRNA-seq (23% of all cells), while snRNA-seq robustly detected this population (87.9-92.8% of all cells) (**Extended Data Fig. 5c**). This is consistent with prior studies of NSCLC using scRNA-seq that showed disproportionally low recovery of malignant cells^6^, suggesting that these cells are vulnerable to sample processing, and emphasizes another potential advantage of snRNA-seq.

To test the feasibility and strengths of these methods, we chose two application cases. First, we obtained sequentially collected (before, and two on-treatment) biopsy specimens from a patient (who a achieved a partial response) treated on the first clinical trial (KEYNOTE-001)^13^ using anti-PD-1 antibody MK-3475 (now known as pembrolizumab) (**Extended Data Fig. 6a**). Although these samples were >10 years old, we achieved excellent technical quality (**Extended Data Fig 6b**) with minimal artifactual gene expression (**Extended Data Fig 6c**), while revealing cellular (**Fig. 2a, Extended Data Fig. 7a,b**) and TCR clonotype diversity. We inferred CNAs and identified distinct clones using k-means clustering (**Fig. 2b, Methods**). Among malignant cells, we noted evolving aneuploidy patterns pre- and on-treatment, suggesting underlying chromosomal instability (**Fig. 2b**), with evidence for immune editing (clones 0 and 3) and immune evasion (clones 1 and 2) of different clones (**Fig. 2c,d**) over time. Interestingly, immune resistant clones 1 and 2 in fact emerged from a small sub-population of pre-existing cancer cells (prior to receiving therapy) and had a temporally conserved cell state strongly enriched for expression of cancer cell intrinsic signatures of immunotherapy resistance (**Fig. 2e**) and de-differentiation (**Fid. 2f**). Importantly, despite the expression of antigen-presentation and IFNg-pathway genes in clone 2 (**Extended Data Fig. 7c**), these cells were strongly enriched for pathways of (de-)differentiation and motility (**Extended Data Fig. 7c,d**), and had increased expression of putative mechanisms of immune evasion, such as *GPX4* (central regulator of ferroptosis)^14^ and *MIF* (**Extended Data Fig. 7c**), and strongly reduced expression of *CD58* (**Fig. 2g**), which was recently identified as an orthogonal mechanism of cancer immune evasion^15^. Among non-malignant cells, we observed increased infiltration with T cells and macrophages (**Fig. 2a, Extended Data Fig. 7a,b**). Integrated analysis of CD8+ T cells revealed infiltration of both stem-like, precursor exhausted, and terminally differentiated cells^16^ (**Fig. 2h**) with corresponding diversification of clonotypes and contraction of pre-existing T cell clones over time (**Fig. 2i**). Together, these findings suggest that, similar to pre-existing resistance mutations which may emerge under the pressure of oncogene-targeted therapies^17^, pre-existing cancer cell clones defined by their underlying CNAs have variable responses to immunotherapies and expand in response to such therapy, *despite* adequate T cell responses.

**Figure 2.**
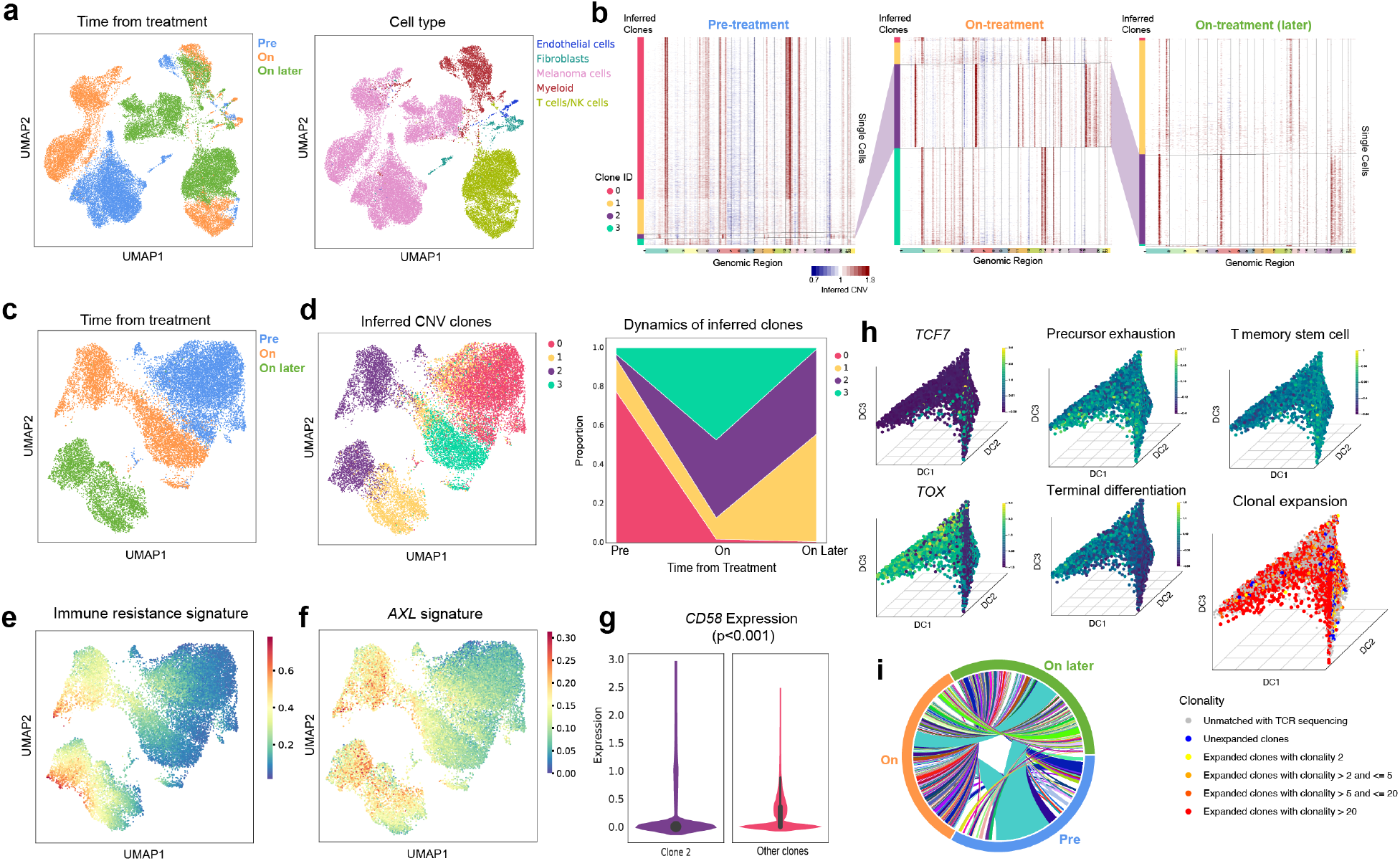
**a**, Merged, unintegrated UMAP and annotation of clusters of cells from sequentially collected specimens during anti-PD-1 therapy. **b**, Inferred copy number alterations across the chromosomal landscape for melanoma cells, in pre- and sequential on-treatment biopsies (from left to right). Genomic location is indicated across the x axis, with chromosomes delineated by vertical lines. Individual cells are plotted along the y axis, with each row representing the CNA profile of one cell, and amplifications in red and deletions in blue. Colored bar to the left indicates clones identified by k-means clustering. **c**, Merged, unintegrated UMAP embedding of cancer cells from three time points indicated by different colors. **d**, (left) Same projection as in (c) indicating cancer clones defined by aneuploidy patterns in (b) and their proportion (right) across pre- and sequential on-treatment biopsies. **e**,**f**, Same projection as in (c) showing expression of (e) immunotherapy resistance program^7^ and (f) AXL-signature from^2^. **g**, Violin plots of expression of *CD58* in emerging clone 2 compared to other cancer clones based on aneuploidy patterns. **h**, Diffusion component (DC) analysis of CD8+ T cells with projections of cells in first 3 DCs colored by indicated genes, signatures and clonotypes. i, Circos plots of T cell receptor clonotypes across different time points (indicated on different aspects of the circle). Connections indicate overlap of identical TCRs between time points.

Second, to demonstrate the scalability of performing such studies in larger cohorts from a multi-institutional clinical trial, we performed snRNA/TCR-seq of 169,015 cells from 20 core needle biopsies collected from 7 patients with liver-metastatic uveal melanoma treated with MEK-inhibitor selumetinib^18^ (**Extended Data Fig. 8a**). Liver metastases occur in most patients with uveal melanoma and are associated with immunotherapy resistance in different cancers, although the underlying mechanisms remain poorly understood^19^. We achieved excellent technical quality (**Extended Data Fig. 8b)** and recovered diverse cell types (**Fig. 3a**), thereby providing a foundational metastatic-niche specific atlas. Among cancer cells, we find that in some patients, rapid changes in aneuploidy patterns (**Fig. 3b**) and distinct transcriptional outputs associated with emerging sub-clones occurred within days of targeted therapy.

**Figure 3.**
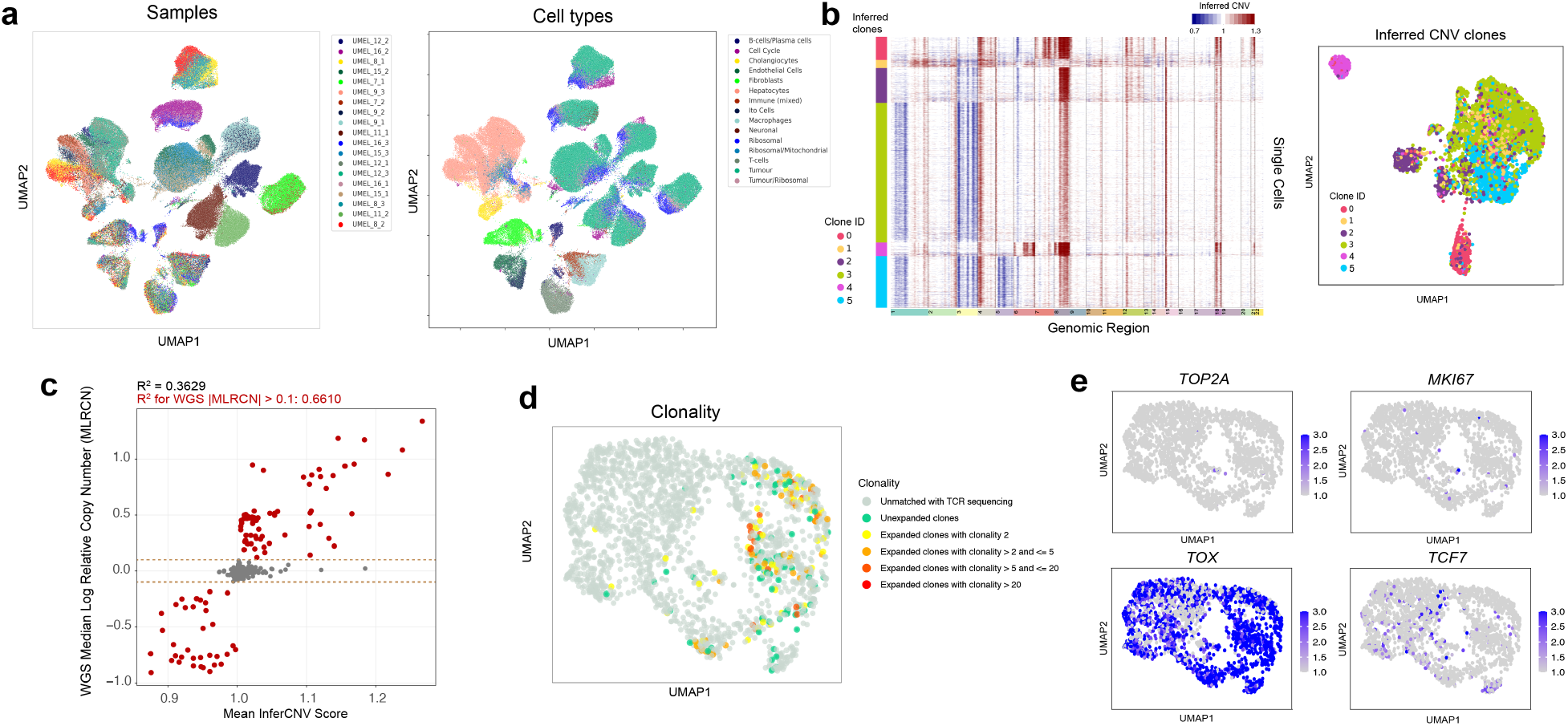
**a**, Merged, unintegrated UMAP and annotated clusters of snRNA-seq transcriptomes across 20 uveal melanoma liver metastasis samples colored by specimen of origin (left) and cell type (right). **b**, Exemplary representation of inferred copy number alterations across the chromosomal landscape of an uveal melanoma liver metastasis specimen (left; color bar on left delineating CNA clones identified by k-means clustering), and corresponding UMAP embedding and clustering (right) colored by respective CNA clones, demonstrating impact of CNA heterogeneity on transcriptional output. **c**, Correlation of chromosome arm copy number alterations predicted by inferCNV from uveal melanoma liver metastases snRNA-seq data vs. population-matched low-pass whole-genome sequencing of the same samples. **d**, UMAP embedding of CD8+ T cells across all uveal melanoma liver metastases with projection of TCR clonotypes. **e**, UMAP embedding (same as in d) of CD8+ T cells across all uveal melanoma liver metastases with projection of selected marker genes of proliferation (*TOP2A, MKI67*) and T cell dysfunction and stemness (*TOX, TCF7*).

Given the apparent importance of aneuploidy patterns in both cohorts presented here, we next established methods to perform simultaneous ulta low pass whole-genome sequencing (ulp-WGS) of the same cell pool on which we performed snRNA-seq (**Extended Data Fig. 9a**). We predicted CNAs in single cell sequencing data using the program inferCNV^2^ (**Extended Data Fig. 9b**), and in the ulp-WGS data using ichorCNA^19^ (**Extended Data Fig. 9c,d**). After integrating the CNA predictions from both sources, we observed a strong correlation between average CNA levels at the chromosome arm level (R2=0.66) (**Fig. 3c**) (**Methods**). This result suggests that single-cell sequencing data is sufficient for robust assessment of single-cell chromosomal instability profiles, and can be used to explore how specific CNAs may impact the corresponding gene expression profiles.

Unlike the tumor microenvironment in other cancer types and metastatic niches, we find that T cells in liver metastases showed very limited clonal expansion (**Fig. 3d**) and were largely composed of dysfunctional T cells with low proliferative capacity (**Fig. 3e**). These results suggest that such cells become progressively dysfunctional or are eliminated within the liver-metastatic milieu and may explain why patients with uveal melanoma have a dramatically lower response rate to ICI compared to cutaneous melanoma.

There are important considerations for future implementation of the approaches outlined here. snRNA-seq performs an unselected detection of cells of the tumor-ecosystem, including capture of cells that are poorly represented in scRNA-seq (e.g. lung cancer cells). Representation of TCRs will be dictated by the *in-situ* fraction of T cell abundances (or lack thereof). However, in frozen tissues, sorting of cell nuclei based on size and scatter patterns may be used to enrich T cells and enhance recovery of matching TCRs. Furthermore, reference atlases are increasingly becoming more helpful in identifying cell types and cell states, such as progenitor-like or exhausted T cells. As snRNA-seq is being more systematically implemented in clinical studies, it will be critical to build such references from frozen tissue specimens to enable rapid definition of cell types and cell states relevant to specific therapies. Lastly, we expect that capabilities for incorporating additional single-cell measurements from frozen tissue (e.g., chromatin accessibility, metabolomics, spatial profiling) will be feasible, and help determine which analytes are best suited to guide clinical application.

In summary, we show that high-quality single-cell transcriptome/TCR profiling and WGS is possible from small, routinely collected clinical specimens. This enables application of these methods to multi-institutional efforts through harmonized and scalable pre-analytical processes while reducing pre-analytical biases, thus representing an important step towards implementing these technologies in clinical care.

## EXTENDED DATA FIGURES AND LEGENDS

**Extended Data Fig. 1.**
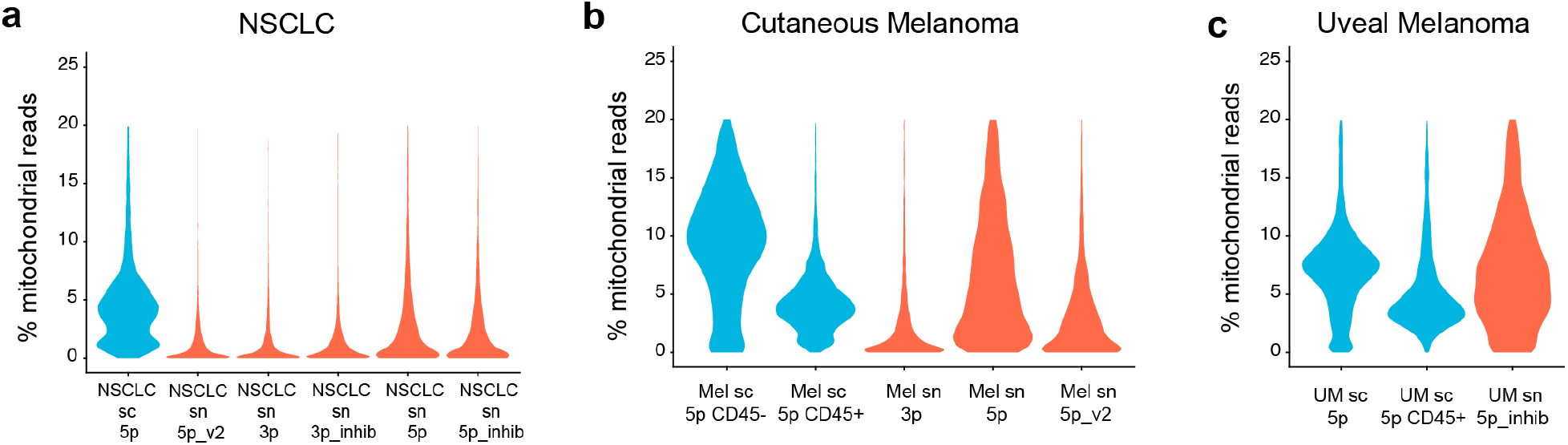
**a-c**, Violin plots indicating percent of mitochondrial reads across samples and different experimental settings in (a) NSCLC, (b) cutaneous melanoma and (c) uveal melanoma. Blue violins indicate scRNA-seq and red violins indicate snRNA-seq.

**Extended Data Fig. 2.**
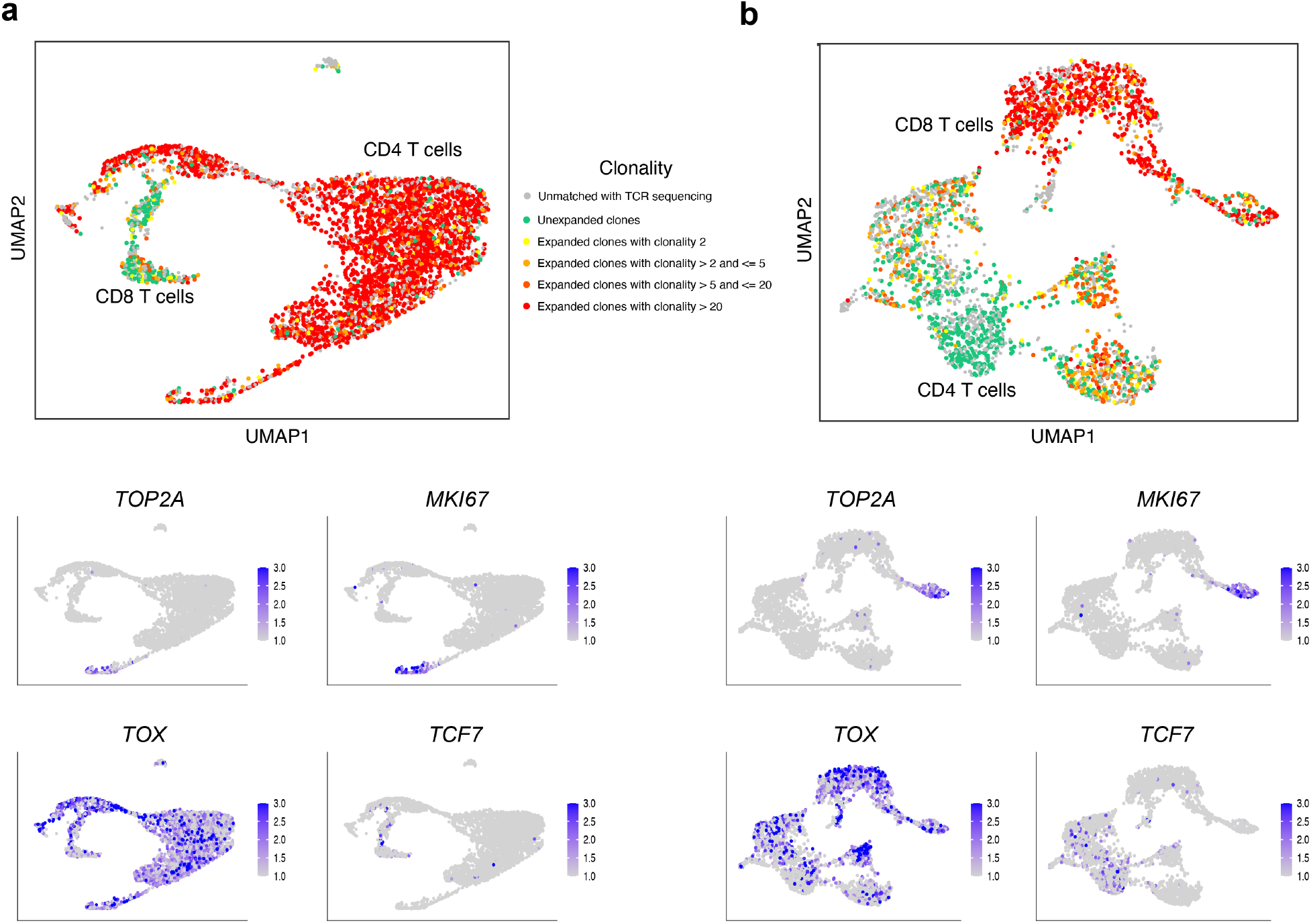
**a**, UMAP clustering of T cells, with projected clonality (top), cell cycle markers and T cell dysfunction and stemness markers (bottom) in primary uveal melanoma. **b**, same as (a) for cutaneous melanoma.

**Extended Data Fig. 3.**
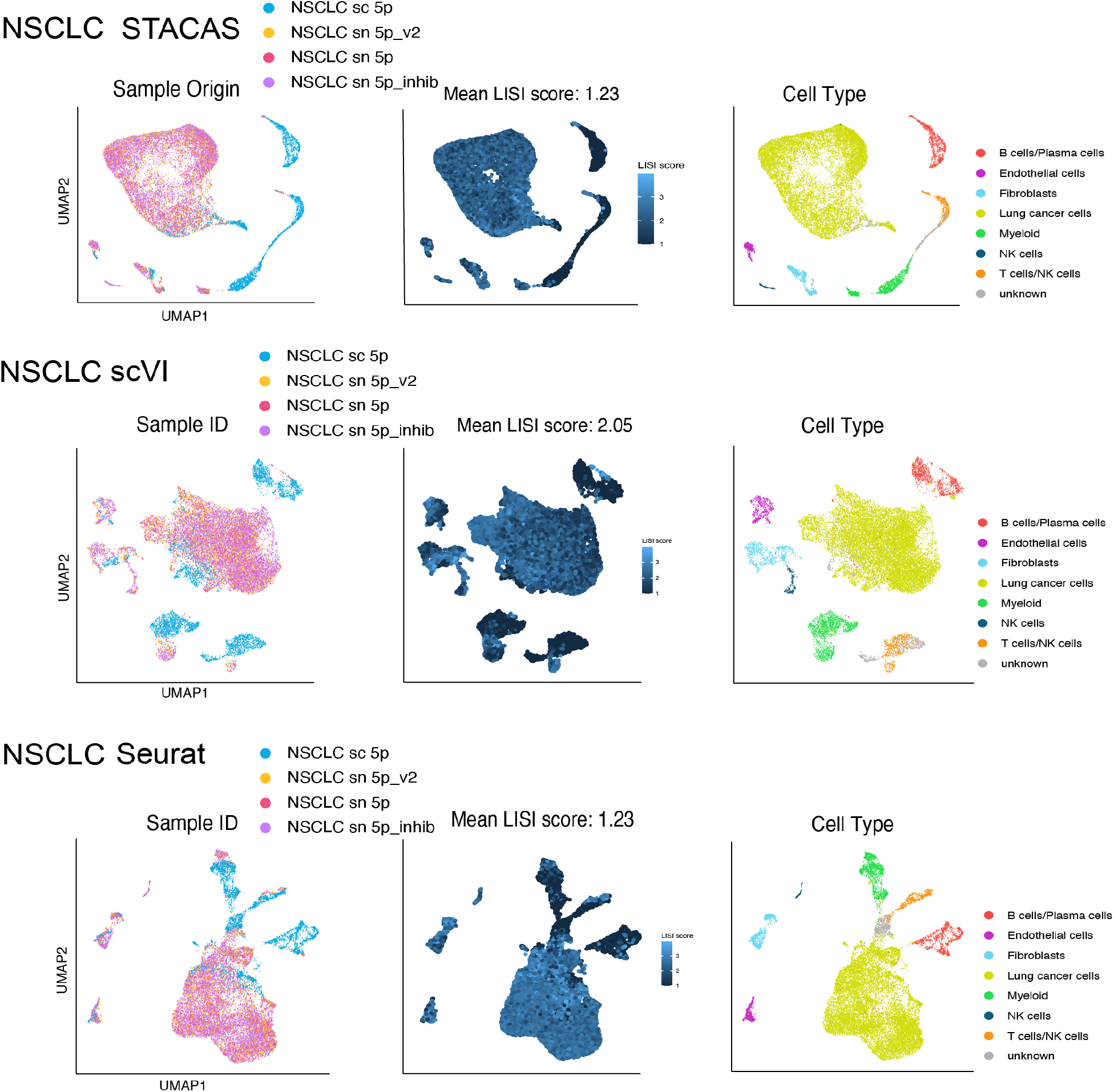
Application of three integration methods in NSCLC.

**Extended Data Fig. 4.**
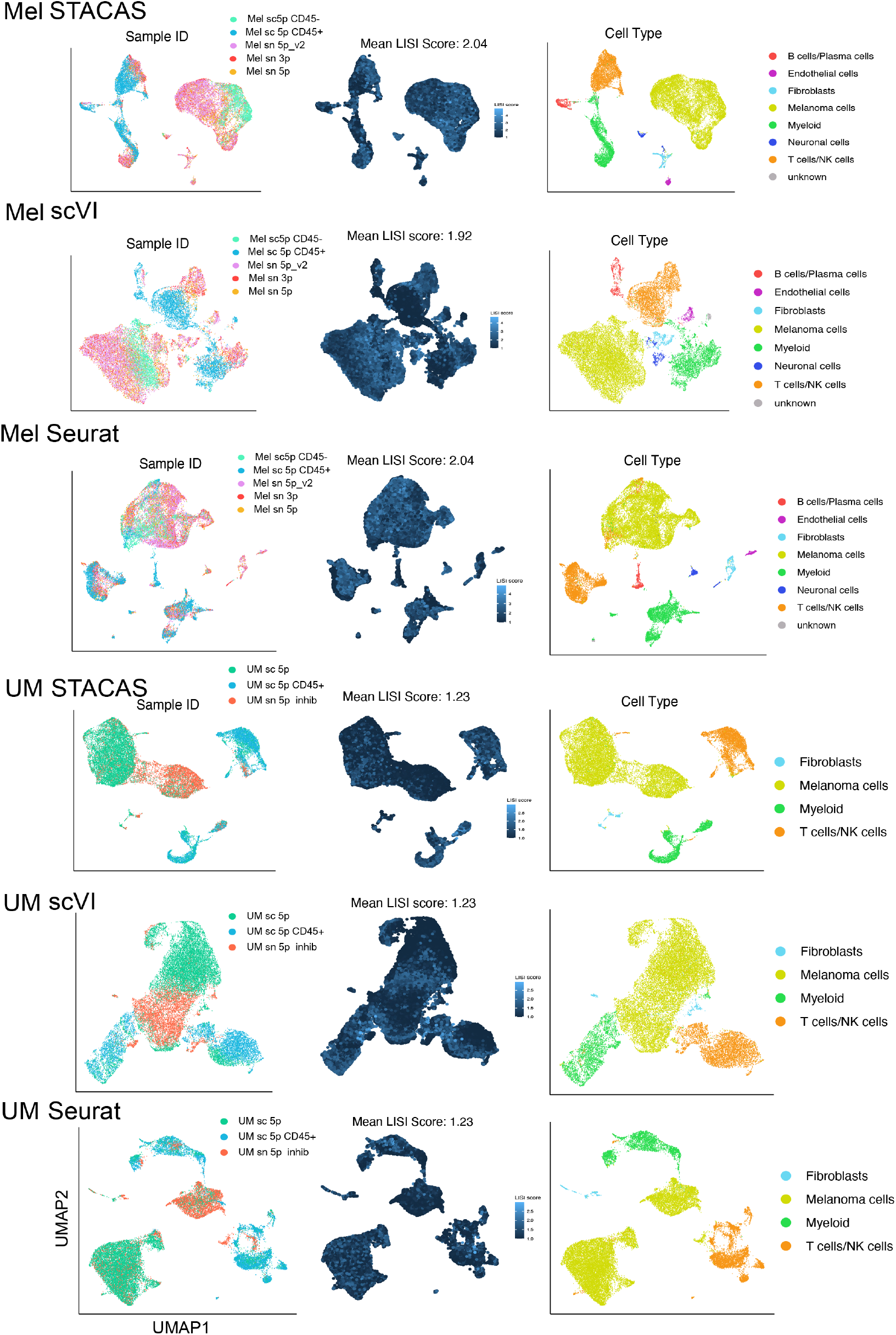
Application of three integration methods across cutaneous and uveal melanoma.

**Extended Data Fig. 5.**
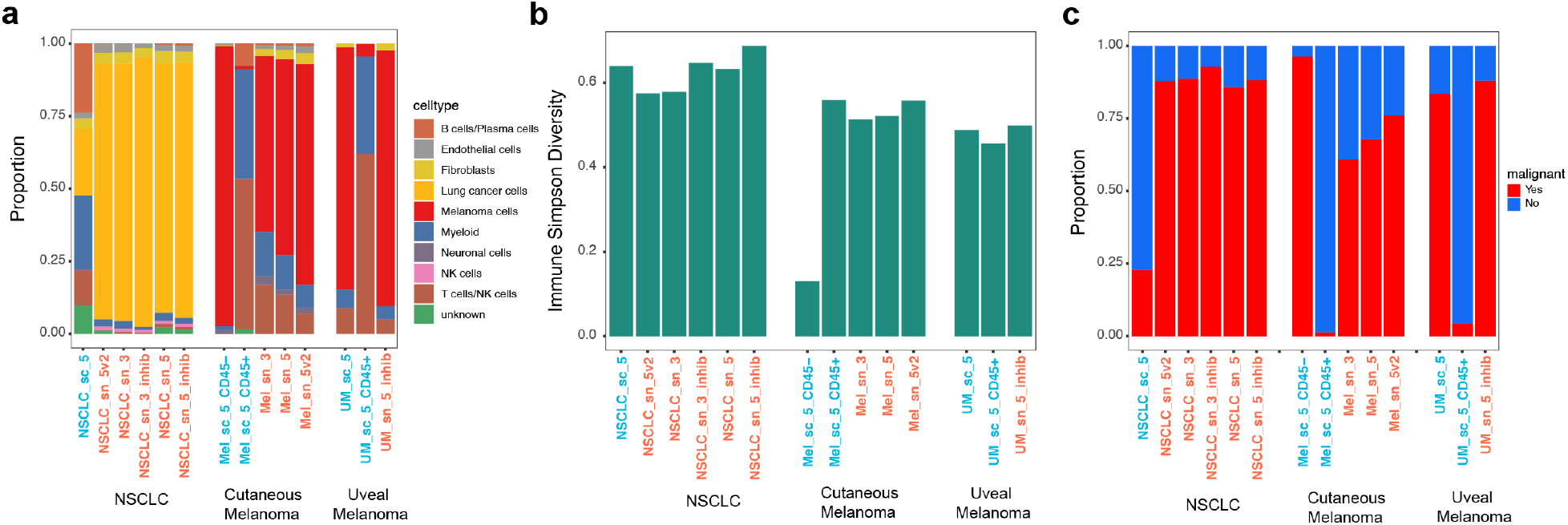
**a**, Stacked bar plots indicating proportions of all cell types in NSCLC, cutaneous and uveal melanoma across different methods. **b**, Simpson diversity index for immune cells in the same samples as (a). **c**, Stacked bar plots of malignant and non-malignant cell fractions in the same samples as (a).

**Extended Data Fig. 6.**
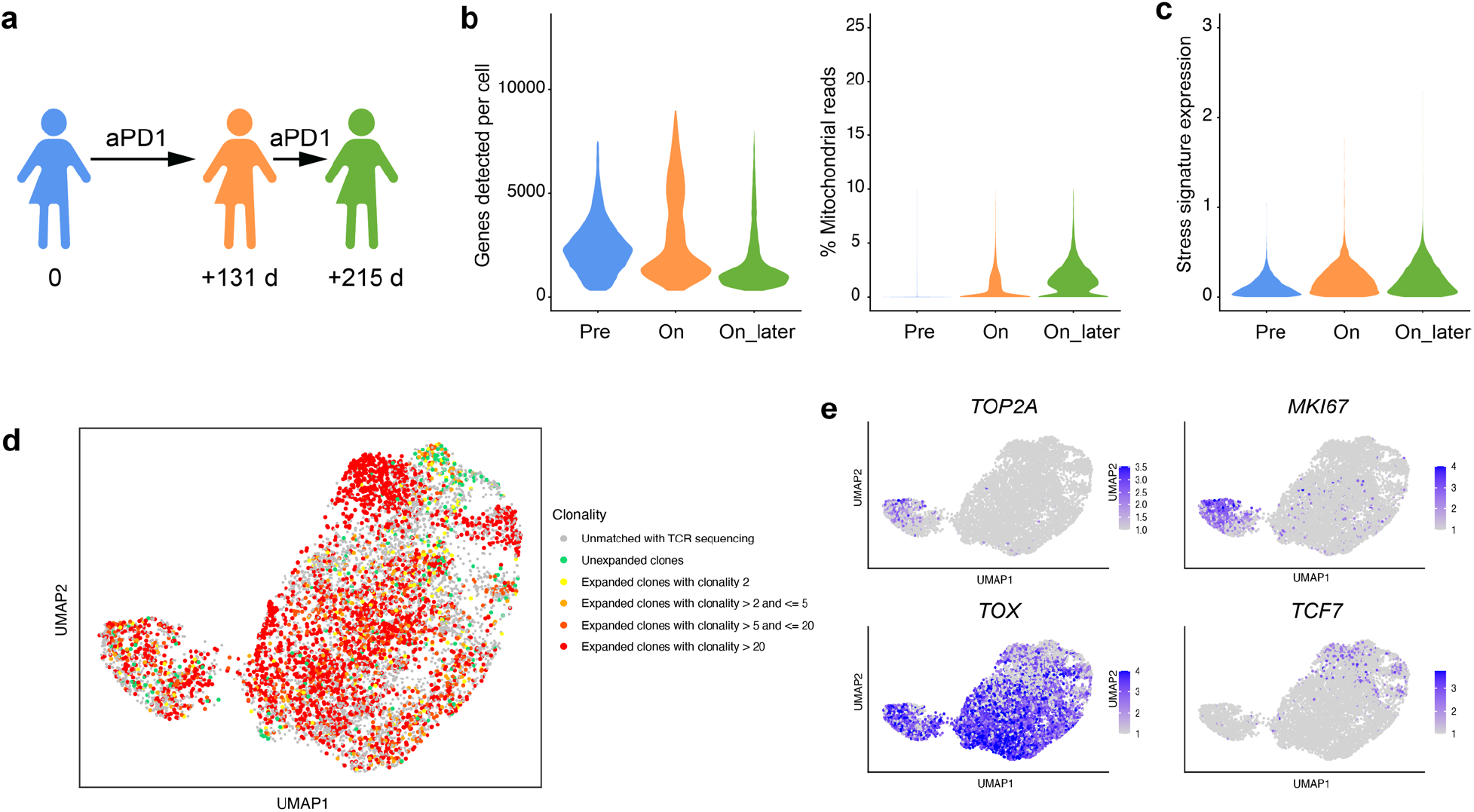
**a**, Timing of sequentially collected specimens in a patient on anti-PD1 therapy. **b**,**c**, Violin plots of (b) genes per cell detected (left) and percent of mitochondrial reads (right), and (c) expression of artifactual signature across samples collected over different time points. **d**,**e**, UMAP representation of CD8+ T cells across all time points and with projected TCR clonality, and (e) cell cycle markers (top) and T cell dysfunction and stemness markers (bottom).

**Extended Data Fig. 7.**
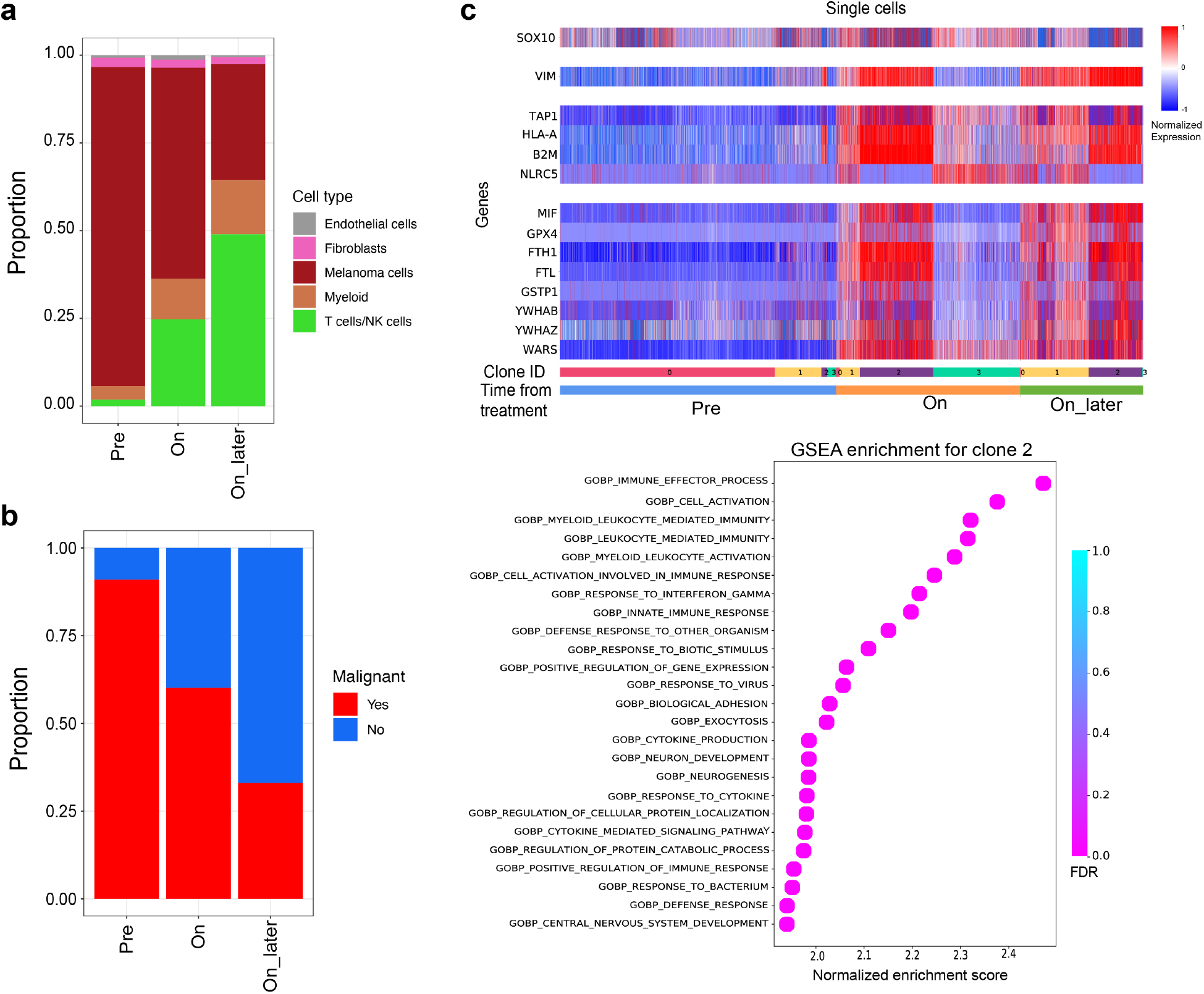
**a**,**b**, Stacked bar plots indicating proportion of (a) all cell types across sequentially collected anti-PD1 therapy tissue specimens and (b) malignant and non-malignant fractions. **c**, Heatmap of selected genes (rows) and their gene expression (normalized expression) in individual cells (column). Indicated on the bottom are time points of sample collection (pre, on, on_later) and clones (0-3) as defined in Fig. 2. **d**, Selected Pathways significantly enriched in Clone 2.

**Extended Data Fig. 8.**
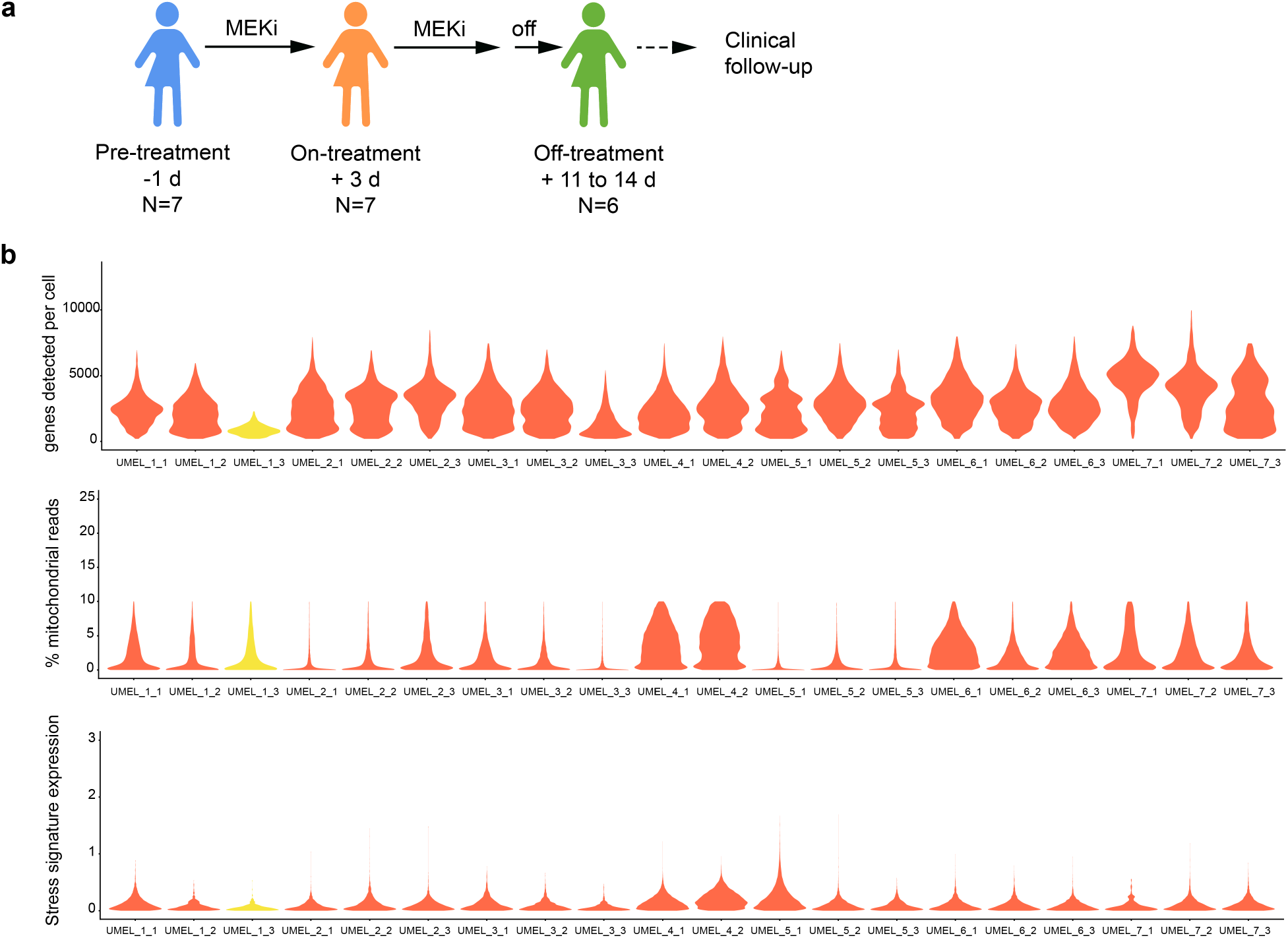
**a**, Schematic of tissue collection and indicated number of specimens per time point. MEKi, MEK-inhibitor (Selumetinib). **b**, Violin plots indicating number of genes detected per cell (top lane), percent of mitochondrial reads (middle lane) and expression of a stress signature (bottom lane) across 20 uveal melanoma specimens. Yellow color indicates a specimen sequenced with 10X 3’ chemistry, with lower quality, while data for the remainder of samples (indicated in red) were generated with 5’ chemistry.

**Extended Data Fig. 9.**
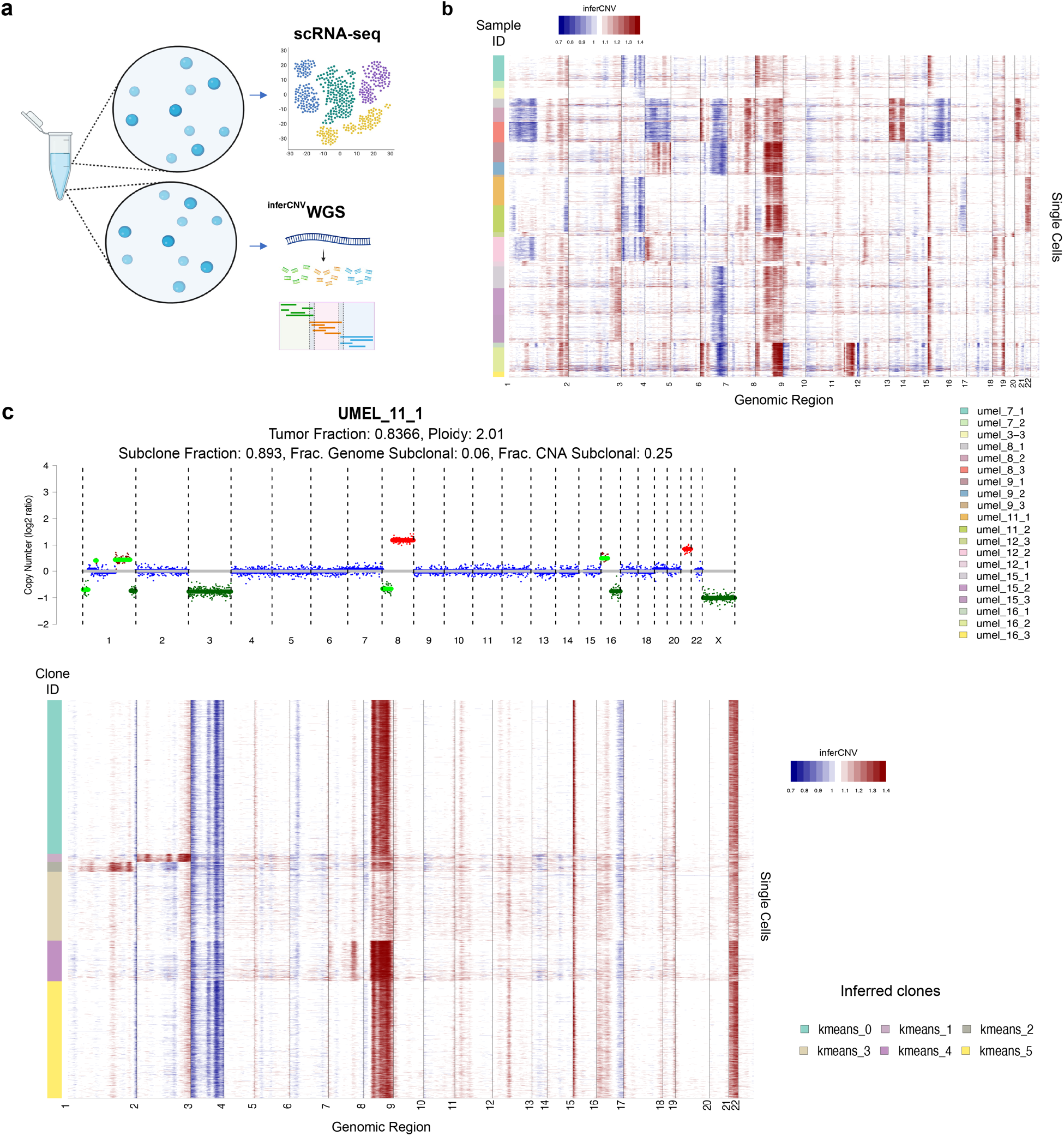
**a**, Schematic design of generation of (low-pass) whole-genome sequencing from the same cell/nucleus pool that was also used for single-nucleus RNA and TCR sequencing. **b**, Inferred CNAs (columns) across samples (indicated by bar on the left) in the uveal melanoma cohort. **c**, Exemplary whole-genome sequencing result (top) showing copy number alterations (y axis, log2 ratio) with amplifications in red, deletions in green and unaltered chromosome regions in blue. Inference of CNAs of the using snRNA-seq that was generated from the same starting cell/nucleus pool as WGS.

## METHODS

### Patient tissue collections

Fresh and frozen tissue specimens were collected under IRB approved protocols at New York Presbyterian Hospital/Columbia University Medical Center (AAAT7416, AAAT2278), Dana Farber Cancer Institute, and University of California, Los Angeles. Surgical specimens (Melanoma brain metastasis, non-small cell lung cancer brain metastasis and primary uveal melanoma specimens) were allocated by qualified pathologists according to institutional guidelines and immediately placed in ice-cold RPMI 1640 (Thermo Fisher, #21875034) without supplements and transported to the laboratory space for immediate processing using single-cell RNA-sequencing and parallel collection of matched flash frozen specimens. Frozen uveal melanoma liver metastases were collected as fine-needle aspiration biopsies during a trial of targeted MEK inhibition^18^. Frozen sequential biopsies of cutaneous melanoma prior and during treatment with anti-PD-1 therapy were collected during the KEYNOTE-001 trial^13^. All procedures performed on patient samples were in accordance with the ethical standards of the IRB and the Helsinki Declaration and its later amendments.

### Fresh tissue specimen processing

All steps but the digestion were carried out on wet ice with pre-cooled buffers. Tissue specimens were weighted and split in half, and pieces of ∼5-8 mm edge length were placed in cryovials and snap frozen in liquid nitrogen before storage at −80°C. The remaining tissue was kept in ice cold RPMI in a petri dish and cut into 1 mm^3^ cubes using two scalpels. The cubes and RPMI were transferred to a 50 ml Falcon tube (Corning) using a 10 ml serological pipette and collected by 5 min centrifugation at 300 x g at 4°C. The tissue was then digested using human tumor dissociation enzymes (Miltenyi Human Tumor Dissociation Kit, #30-095-929) according to manufacturer instructions based on the tissue weight. Briefly, tissue was resuspended in pre-warmed RPMI and human tumor dissociation enzymes were added. The sample was then placed in a 37°C water bath and agitated every 2 minutes. Every 5 minutes the tissue was further mechanically dissociated by pipetting using pipettes of decreasing orifice. This process was continued until most of the tissue had dissociated. After 10 minutes (Melanoma) or 15 minutes (Lung cancer) the samples were filtered through a pre-wetted 70 µm cell strainer (Corning) into a new 50 ml Falcon tube, collected by centrifugation for 5 min at 400 x g and 4°C, and the supernatant was decanted. The cell pellet was resuspended in 3 ml ACK buffer (Thermo Fisher; #A1049201) to lyse red blood cells. After 1 min incubation the reaction was terminated by dilution with 30 ml ice cold sorting buffer (2% Fetal bovine serum/1mM EDTA in PBS). The cells were collected by centrifugation for 5 min at 400 x g and 4°C, resuspended in 1 ml ice cold sorting buffer and cell count and viability was assessed using trypan blue and disposable Neubauer counting chambers (Bulldog Bio, Inc. Portsmouth, NH). Cells were then allocated for direct loading (non-small cell lung cancer and primary uveal melanoma), or further processed for fluorescence-activated cell sorting (Cutaneous melanoma and primary uveal melanoma).

### Fluorescence-activated cell sorting

To enrich viable immune cells (primary uveal melanoma sample) or sort viable immune and non-immune cells (cutaneous melanoma) the samples were sorted using a FACS Aria II (BD Biosciences). First, cells were stained for viability (Zombie NIR, 1:500 in PBS; Biolegend, San Diego, Ca; #423106) for 10 min at room temperature in the dark. Thereafter, cells were washed once with sorting buffer, collected by centrifugation and surface antigens were stained for 15 minutes on ice in the dark. The primary uveal melanoma sample was stained with Pacific-Blue-aCD45 (Biolegend, #304022). The cutaneous melanoma sample was stained with the following (all Biolegened): Human TruStain FcX (#422302), Pacific-Blue-aCD45 (#304022), PE-Dazzle594-aCD3 (#300450), PE-CY7-aCD66b (#305116), APC-aCD15 (#301908). After staining, the samples were washed twice with ice-cold sorting buffer and 1.5×10^3^ cells per population of interest were sorted and immediately processed for scRNA-sequencing.

### Single cell RNA library preparation

Sorted and unsorted single cell suspensions (1.2-1.5×10^3^ cells) were transferred into low-binding 1.5 ml Eppendorf tubes (Eppendorf, Hamburg, Germany), centrifuged and washed twice with 1 ml loading buffer (PBS with 0.05% RNase-free BSA; Thermo Fisher, #AM2616) using a swinging bucket centrifuge at 4°C with 400 x g for 5 min. After the final spin all but 31 µl buffer were removed and the samples were loaded on a Chromium controller with Chromium Single Cell V(D)J Reagents (10x genomics, Pleasanton, CA; #1000006) for 5’ RNA capture. Gene expression libraries were then generated using Chromium Single Cell 5’ Library construction kit (#1000020) according to manufacturer instructions.

### Single-nuclei extraction from tiny frozen specimens

We adopted the previously described salt-tris (ST) based extraction method^9^ and implemented critical changes to enable tissue sparing extraction of nuclei for single-nuclei RNA-sequencing from clinical-grade frozen tissue specimens and leverage minute specimens such as fine-needle aspiration biopsies (FNAs). To this end, we used a Leica CM1950 cryostat (Leica, Wetzlar, Germany) for initial tissue processing and additional washing steps for OCT removal. Frozen tissue specimens with 2-10 mm edge length were embedded in optimal cutting temperature (OCT) compound (Tissue-Tek, Sankura) on dry ice. Samples were then mounted on sample holders and excess OCT was trimmed away using a blade leaving ∼5 mm OCT around each side of the tissue. Fine needle aspiration biopsies were directly mounted on sample holders of the cryostat using a small amount of OCT. Multiple 20µm tissue curls were cut per tissue and collected in pre-cooled 5 ml tubes (Eppendorf, Hamburg, Germany) ensuring no thawing while transferring and stored on dry ice until processing. The number of curls required depends on the tissue size and ranges from 3-4 curls for large specimens (1 cm^2^) to 10-15 for FNA’s. All subsequent steps were performed on wet ice and all centrifuges were equipped with swinging buckets and cooled to 4°C. For extraction of nuclei, the tubes were moved from dry ice to wet ice and left to equilibrate briefly. After 30 seconds, 4 ml of ice-cold PBS without calcium or magnesium (Thermo Fisher) were added and the tubes were inverted until all OCT had dissolved and the clean tissue could be collected by centrifugation at 300 x g for 2 min. The tissue was then resuspended in 1 ml ST buffer [146 mM NaCl, 10 mM Tris-HCL pH7.5, 1mM CaCl2, and 21 mM MgCl2 in ultrapure water] with 0.03% Tween-20 Sigma Aldrich, p-7949), 0.1% BSA (New England Biolabs, B9000S) (TST buffer) and supplemented with or without 40 U/ml RNAse inhibitor (RNAse OUT, Thermo Fisher) (**Extended Data Table 1)**. The suspension was thoroughly pipetted 15 x using a 1 ml pipette to mechanically dissociate the tissue and left to incubate for 5 min on ice. After 5 min the pipetting step was repeated, and the reaction was quenched using 4 ml ST buffer with or without 40 U/ml RNAse inhibitor (RNAse OUT, Thermo Fisher) (**Extended Data Table 1**). The sample was filtered through a pre-wetted 70 µm nylon mesh filter (Fisher Scientific) into a 50 ml conical tube and the filter was washed with 5 ml ST buffer. The tube was then centrifuged at 500 x g for 5 min to collect the dissociated nuclei. After carefully decanting the supernatant, the nuclei were resuspended in 100-400 µl ST buffer without RNAse inhibitor and filtered with a 40 µm mesh filter attached to a FACS tube (Fisher Scientific). The nuclei concentration and dissociation quality was then determined in a 5 µl aliquot using Neubauer counting chambers (Bulldog Bio, Inc. Portsmouth, NH) and a fluorescent microscope (EVOS FL, Thermo Fisher) after staining nuclear DNA with 50µg/ml Hoechst 33342 (Thermo Fisher).

### Single nuclei RNA library preparation

0.9-1.5×10^3^ nuclei were loaded in ST buffer without RNAse inhibitor using a Chromium controller and chromium reagents (10x genomics) for 3’ or 5’ capture as indicated (**Extended Data Table 1**). After reverse transcription and cleanup, cDNA libraries were generated according to manufacturer instructions with one additional cycle of cDNA amplification to account for the relatively lower amount of RNA in nuclei compared to whole cells.

### Single cell and nuclei TCR library preparation

Single cell and single nuclei TCR libraries were prepared from amplified cDNA libraries according to manufacturer instructions using the following reagents (all 10x genomics): Chromium Single Cell V(D)J Enrichment Kit for human T cells (#1000005) was used for cDNA generated with Chromium Single Cell V(D)J reagents (#1000006), and final sequencing libraries were prepared using Chromium i7 multiplexing kit (#120262) Single Cell Human TCR Amplification Kit (#1000252) was used for cDNA generated with Chromium Next GEM Single Cell 5’ v2 reagents (#1000263), and final sequencing libraries were prepared using Library construction kit (#1000190) and Dual Index Kit TT set A (#1000215).

### Sequencing of single cell and single nuclei libraries

Final sequencing libraries were quantified using Tapestation D1000 and D5000 reagents (Agilent) and a 2200 TapeStation system. Samples were then mixed and sequenced to target >20,000 reads per cell for gene expression libraries and >5,000 reads per cell for TCR libraries using NovaSeq S4 or HiSeq 4000 (Illumina. San Diego, CA) with at least 2×100 BP coverage (**Extended Data Table 1**).

### Genomic DNA extraction for low-pass whole genome sequencing

Excess nuclei (>1×10^5^) from the sample preparations for sn-RNAseq were collected by centrifugation (500 x g, 5 min) and snap frozen after removing all but ∼10 µL ST buffer and stored until further processing at −20°C. If insufficient numbers of nuclei were available after loading, additional curls were processed using the same methods as described above for single nuclei extraction. To extract genomic DNA from nuclei the nuclei were briefly thawed on wet ice and genomic DNA was extracted using DNAeasy Blood and Tissue kit (Qiagen, Hilden, Germany) according to manufacturer instructions and eluted in RNAse and DNAse free water at 37°C for 5 minutes. The DNA concentration was then quantified using a Nanodrop.

### Library construction for ultra low pass whole genome sequencing

20-50 ng genomic DNA served as input for ultra low pass whole genome sequencing (ulp-WGS) library preparation with the Lotus DNA Library Prep Kit (Integrated DNA Technologies - IDT, Coralville, IA) using xGen™ Stubby Adapter (IDT) and UDI Primer Pairs (IDT) according to manufacturer institutions with the following specifications: In the enzymatic preparation program, samples were held at 32°C for 9 minutes. Adapters were not diluted in the ligation step and the optional PCR (5 cycles) and cleanup step were performed so the stubby adapters could be amplified with primers to incorporate the index sequences and P5 and P7 sequences. After library cleanup with AMPure beads (Beckman Coulter, Brea, CA), the libraries were quantified with Qubit 1X dsDNA HS Assay Kit (ThermoFisher) and Tapestation D5000 HS tapes (Agilent) and stored at −20°C until sequencing.

### ulp-WGS sequencing and copy number assignment

Indexed WGS-libraries were mixed equimolarly and sequenced on an Illumina MiSeq instrument with 0.1X coverage using the V2-300 cycle kit (Illumina). Using Illumina pipelines, .bam and .bai files were generated from .fastq files which served as input for ichorCNA^20^ generating .seg files for visualization. Finally, GISTIC 2.0^21^ was used to assign a copy number to each gene.

### Processing FASTQ files into gene expression matrices

Demultiplexed FASTQ files from raw single-cell RNA sequencing reads were aligned to the human GRCh38 genome, and gene counts were quantified using Cell Ranger ‘count’ (v5.1 for uveal melanoma liver metastasis and sequential anti-PD1 therapy samples; v6.1 for all other datasets; 10x Genomics). Reads mapping to introns and exons were counted in order to increase the number of genes detected per nucleus and number of nuclei passing quality control, leading later on to improved cell -type identification. To include introns during read mapping, we followed the instructions provided by 10x Genomics (https://support.10xgenomics.com/single-cell-gene-expression/software/pipelines/latest/advanced/references), and created a ‘pre-mRNA’ human GRCh38 reference genome, using the Cell Ranger command ‘mkref’ with a customized gene transfer format (GTF) file, in which all transcripts were labelled as exons.

### Removal of background noise in gene expression matrices

We used the ‘remove-background’ function of CellBender (v0.2.0) on CellRanger-generated ‘raw_feature_bc_matrix.h5’ files, in order to remove technical ambient-RNA counts and empty droplets from the gene expression matrices^22^. The parameter ‘expected-cells’ was obtained from the Cell Ranger metric ‘Estimated Number of Cells’, while the parameter ‘total-droplets-included’ was set to the midpoint of a plateau in the barcode-rank plot, based on visual inspection.

### Quality control and filtering

After processing by CellBender, expression matrices were processed individually in R (v4.0.2) using Seurat^23^ (v4.0.1). For sequentially collected specimens from the patient on anti-PD1 therapy, we applied filters to keep only cells with 300-10000 genes, 400-30000 UMIs, and <10% of mitochondrial reads. For uveal melanoma liver metastasis samples, we kept cells with at least 200 genes, 100 UMIs, and <10% of mitochondrial reads. Additionally, we removed cells with gene or UMI counts that were above a sample-specific maximum limit, based on the observed saturation curve between gene and UMI counts for each sample. Specifically, we visually inspected each of these curves, and observed the point at which outlier cells, in terms of very large gene and/or UMI counts, started to appear. We set the maximum limits so as to exclude these outliers for each sample, and the list of these limits is given in Extended Data Table 4. Finally, for all other datasets, we kept cells with at least 300 genes and <20% of mitochondrial reads. Additionally, Scrublet^24^ expected doublet rate of 9.6%.

Filtered gene-barcode matrices were then normalized with the ‘NormalizeData’ function using the ‘LogNormalize’ method. The top 2,000 variable genes were identified using the ‘vst’ method in the ‘FindVariableFeatures’ function. Gene expression matrices were scaled and centered using ‘ScaleData’. We performed principal component analysis (PCA) as well as uniform manifold approximation and projection (UMAP) dimension reduction using the top 30 principal components. We also used the AddModuleScore function of Seurat to calculate the expression of a stress-associated gene signature (Extended Data Table 5).

### Manual cell-type annotation

We integrated and clustered samples within each dataset using Seurat, and identified differential gene expression (DGE) between clusters using the FindAllMarkers function. We then manually annotated cell type clusters using the following list of marker genes (**Extended Data Table 5**), except for B-cells/Plasma cells, which were annotated on the basis of IG genes being differentially upregulated in them. This initial labeling resulted in the identification of major cell types such as melanoma cells, lung cancer cells, hepatocytes, T cells/NK cells, and others (**Extended Data Table 5**). Next, to annotate subtypes of T cells/NK cells only, we created a Seurat object containing only these cells, and reran scaling, PCA, UMAP dimension reduction, clustering and DGE analysis. The resulting clusters were again annotated manually as CD4+ T-cells, CD8+ cells, T-regs and others (**Extended Data Table 5**).

### Copy Number Alteration (CNA) Inference Using InferCNV

Chromosomal CNA profiles of individual cells were inferred from transcriptional data using inferCNV^2^ (v1.6.0). For each sample, we used cells that were identified as immune cells by SingleR^25^ as a diploid reference to estimate CNAs in the non-immune cells. We applied a cutoff of 0.1 for the minimum average read counts per gene among reference cells/nuclei, set the clustering to ‘subcluster’, denoised the output using the default ‘sd_amplifier’ of 1.5, and ran Hidden Markov Models (HMM) to predict the CNA level.

### Arm-level inferred CNA comparison between snRNA and scRNA samples

To measure correlation of cancer cell inferCNV profiles between snRNA and scRNA-seq samples (**Fig. 1i**), for datasets that included both types of samples, we calculated average inferCNV scores for each chromosome arm for the cancer cells in each individual sample. To exclude chromosome arms that did not exhibit any large-scale amplifications or deletions, we filtered out arms that had an average inferCNV score between -.01 and .01. We also did not consider the CD45+ sample from our melanoma brain metastasis dataset, as it mostly contained immune cells, or the NSCLC 3’ sequencing protocols, as these appeared to be of lower quality in terms of gene counts and stress signature expression than the 5’ protocols (**Fig. 1d**). For every possible pair of one fresh vs. one frozen sample in each of our datasets, we then calculated the Spearman correlation between average chromosome arm inferCNV scores in each of the two samples.

### TCR data processing and integration

TCR FASTQ files were aligned using Cell Ranger ‘vdj’ (v6.1.1; 10x Genomics). We then used the combineTCR function the package scRepertoire to process filtered contig annotation files from the Cell Ranger output. Clonotypes which were annotated as having the same clonotype gene by combineTCR, and occurred in two or more samples within a dataset, were considered as shared within those samples.

### Comparison of TCR clonotype composition between fresh and frozen samples

For our primary uveal melanoma and cutaneous melanoma brain metastasis TCR datasets (**Fig. 1g,h**), we used a hypergeometric test to compare clonotype compositions between every pair of fresh vs. frozen samples. We used the clonotype composition of the fresh sample as a reference, and compared it with the composition of the frozen sample. Specifically, we used the clonotypes that were shared between the fresh sample and frozen sample as input to the x argument of the dhyper function in R, the clonotypes found in the fresh sample as the m argument, clonotypes found in the frozen sample but not the fresh sample as the n argument, and the total number of clonotypes in the frozen sample as the k argument.

For each sample, we also calculated the Gini coefficient of the distribution of TCR clonotype frequencies, using the gini function downloaded from https://github.com/oliviaguest/gini/blob/master/gini.py

### Comparison of arm-level CNAs between single-cell inferCNV predictions and ulp-WGS measurements in uveal melanoma liver metastases

Using Illumina pipelines (automatic on miSeq machine), .bam and .bai files were generated from each sample from .fastq. ichorCNA^20^ analysis of these files generated .seg files for visualization. GISTIC 2.0^21^ was used to assign a copy number to each gene. We calculated average inferCNV scores for each chromosome arm for the cancer cells in each sample of our uveal melanoma liver metastases dataset. We then compared this with the median log relative copy number measured for each chromosome arm using ulp-WGS (**Fig. 3c**). We calculated the Spearman correlation between these two values using two settings. First, we included all chromosome arms, and second, to exclude chromosome arms that did not exhibit any large-scale amplifications or deletions, we filtered out arms that had a ulp-WGS median log relative copy number between -.1 and .1.

### Diffusion component (DC) analysis

For CD8+ T-cells in our three sequential anti-PD1 therapy samples (**Fig. 2h**), we computed DCs using the ‘DiffusionMap’ function of the Destiny R-package^26^. The ‘AddModuleScore’ function in Seurat was applied to calculate average expression levels of several T-cell gene signatures on a single-cell level (Extended Data Table 5). We plotted expression of several of these signatures, as well as expression of the TOX and TCF7 genes, and TCR clonotype expansion, on the first three diffusion components.

### Evaluation of batch correction methods on fresh and frozen samples

To determine the extent of integration achievable between samples of fresh and frozen origin, we applied the following set of batch correction methods: STACAS^27^, scVI^28^ and Seurat (**Extended Data Fig. 3,4**). For integration methods, genes were filtered using scanpy by selecting only the top 8000 highly variable genes^29^. Integration results were visualized using UMAP. For scVI, nearest neighbors were computed in the reconstructed gene space (with PCA preprocessing); for STACAS and Seurat, UMAP was computed in the integrated space with PCA preprocessing. The degree of integration achieved by each method was evaluated by computing the LISI score^12^. For all methods, LISI scores were computed on 20 principal components and visualized using UMAP. The mean LISI score was also computed for each method.

### Identification of Differentially Expressed Genes

We applied Scanpy’s function for ranking genes to identify differentially expressed genes. For each cluster (defined by K-means clustering), we compare the genes contained within the cluster to genes not contained in the cluster using a t-test with Benjamini-Hochberg correction. Genes are then ranked by scores, and the top 300 genes for each cluster are returned (excluding mitochondrial and ribosomal genes).

### Analysis of inferCNV clonal dynamics in the KEYNOTE-001 patient

#### Preprocessing

Cells from all treatment time-points were normalized by library size together and log transformed using scanpy.pp.normalize_per_cell and scanpy.pp.log1p^29^ (**Fig. 2a**). Batch correction was not performed, due to the observation that immune cells across samples showed more overlap than tumor cells. We then selected tumor cells from the normalized anndata object from the KEYNOTE-001 patient only. For our data, we verified tumor cell identification using both inference of copy number alterations that were expected to be present in malignant cells using inferCNV^2^ (v1.6.0) (**Fig. 2b**), as well as known lineage marker genes, including *MITF, MLANA*, as well as the MITF-high and AXL-high signature gene sets^2^ (**Fig. 2e,f**).

#### Clustering of inferCNV clones

We utilized K-means clustering on copy number data generated from inferCNV to group cells into clones defined by shared patterns in copy number alterations across genes (**Fig. 2b**). We found through visual inspection of inferCNV results that, across all treatment time points, there appeared to be four distinct groups of cells, each having a unique inferCNV footprint. Thus, K-means clustering was performed with k=4. Visualization via UMAP was utilized to analyze clonal dynamics with treatment. Temporal analysis was performed by plotting the proportion of cells belonging to each clone at each treatment time-point. Expanding clones were then defined as those that showed an increase in proportion at the “on_later” time point compared to the “pre” time point (**Fig. 2d**).

#### Characterization of differentially expressed genes in expanding clones

We selected 300 differentially expressed genes for each clone, via the procedures outlined above. The differentially expressed genes for expanding Clone 2 were analyzed using the preranked option in GSEA^30^ and the c5.goBP curated set of genes. The normalized enrichment score and false discovery rate for the top 25 enriched genesets was visualized (**Extended Data Fig. 7d**).

#### Analysis of patterns exhibited by known genes/gene signatures

Characterization of clonal and treatment-induced dynamics was performed by analyzing the expression of genes belonging to previously defined geneset signatures, including immune checkpoint inhibitor resistance (ICR) signature^7^ and the AXL-high signature^2^. Normalized expression was averaged across all genes belonging to the signature set, and plotted using the UMAP representation (**Fig. 2e,f**). Individual genes of interest were manually selected and visualized using a heatmap, where cells were grouped on the x-axis according to treatment groups and clones within each (**Extended Data Fig. 7c**). For each gene, z-scoring was performed across all cells to normalize the data and show variability across all cells. Analysis of *CD58* expression (**Fig. 2g**) was performed by partitioning data into two groups: cancer cells belonging to Clone 2 and all other cancer cells. Normalized expression of *CD58* was summarized across cells in each group, and significance was assessed using a Mann-Whitney U test^31^.

## Data and code availability statement

Processed data is available on GEO accession number: GSE192402. Raw data will be available on dbGAP: accession number pending. Code will be made publicly available via: https://github.com/IzarLab/sc_sn_RNA_seq

## Acknowledgments

We thank Hanina Hibshoosh at Columbia University and Matan Hofree for fruitful discussions. B.I. is supported by National Institute of Health (NIH) National Cancer Institute (NCI) grants K08CA222663, R37CA258829, U54CA225088, a Burroughs Wellcome Fund Career Award for Medical Scientists, a Velocity Fellows Award, the Louis V. Gerstner, Jr. Scholars Program and a Young Investigator Award by the Melanoma Research Alliance. R.C., E.A. and B.I. are supported by NCI grant R21CA263381 and a Columbia University Research Initiatives in Science & Engineering Award. E.A. was supported by NCI grant R00CA230195. J.L.F. acknowledges support from the Columbia University Van C. Mow fellowship. G.A-R. and A.R. are supported by Parker Institute for Cancer Immunotherapy and NIH grant P01CA168585. AMT is supported by NCI 5K22CA237733-03.This work was supported by NIH/NCI Cancer Center Support Grant P30CA013696 and the Molecular Pathology Shared Resource and its Tissue Bank at Columbia University.

## Author contributions

B.I. conceived of the study. B.I. and E.A. jointly provided overall supervision of the study. J.C.M., A.D.A., Y.G., P.H., L.C. and S.B. performed experiments. Y.W., J.L.F., J.C.M., Y.G. S.T., G.A-R., A.M.T. performed analyses. S.A.K., B.S.H., A.R., G.K.S. and R.D.C. provided clinical specimens. A.R., R.D.C. and A.M.T. provided additional supervision. Y.W., J.L.F, E.A. and B.I. wrote the manuscript. All authors reviewed, contributed to, and approved of the manuscript.

